# Food web networks shift across a precipitation gradient due to changes in community composition and species interactions

**DOI:** 10.1101/353920

**Authors:** Laura Melissa Guzman, Bram Vanschoenwinkel, Vinicius F. Farjalla, Anita Poon, Diane Srivastava

**Affiliations:** Department of Zoology, University of British Columbia, Vancouver, British Columbia, V6T1Z4, Canada; Community Ecology Laboratory, Vrije Universiteit Brussel, Ixelles, 1050, Belgium; Department of Ecology, Universidade Federal do Rio de Janeiro, Rio de Janeiro, 21941-590, Brazil

**Keywords:** Community composition, environmental gradients, phytotelm macroinvertebrate communities, markov networks

## Abstract

Ecological networks change across spatial and environmental gradients due to (i) changes in species composition or (ii) changes in the frequency or strength of interactions. Here we use the communities of aquatic invertebrates inhabiting clusters of bromeliad phytotelms along the Brazilian coast as a model system for examining turnover in the properties of ecological networks. We first document the variation in the species pools of sites across a geographical climate gradient. Using the same sites, we also explored the geographic variation in species interaction strength using a newly developed Markov network approach. We found that community composition differed along a gradient of water volume within bromeliads due to the turnover of some species. From the Markov network analysis, we found that the top-down effects of certain predators differed geographically, which could also be explained by geographic differences in bromeliad water volumes. Overall, this study illustrates how a network can change across an environmental gradient through both changes in both species and their interactions.

## Introduction

Ecological networks can change across spatial and environmental gradients in two main ways: the composition of species can change, or the strength of interactions between species can change (Tylianakis and Morris 2017a). Species composition can vary across an environmental gradient if the environment filters particular traits (M. A. Leibold et al. 2004; Kraft et al. 2014). Species composition can change across space if species differ in their dispersal abilities (M. A. Leibold et al. 2004; Kraft et al. 2014). Even when species are found together across a gradient, the strength of interactions between these species can vary between sites on the gradient. For example, consumers may find prey more efficiently in structurally simple, as opposed to complex, habitats (Laliberté and Tylianakis 2010). Consumption rates can also be higher in warmer sites, due to temperature-dependence of metabolic rates (Rall et al. 2012). Here we combine multiple analyses to show how environmental and spatial gradients affect both the composition and interactions of species in a ecological network.

Estimating changes between sites in species interaction strengths is notoriously difficult, much more so than estimating changes in community composition (e.g. Anderson 2001; Borcard, Gillet, and Legendre 2011). For example, pairwise competition experiments consider interactions between only two species, ignoring other interactions. Yet, pairwise interactions in real communities may be influenced by other species (Maser, Guichard, and McCann 2007). Experiments that have measured interactions in a community context often remove one species from the system and assess the impact on the whole community Yet, his approach cannot reconstruct the strength of the interactions between all members of a community (Paine 1966). Inferring species interactions from observational data has the advantage of realism. For instance, combining observations of prey abundance and predator foraging rates can provide information on interaction strengths (Wootton 1997). This method, however, cannot estimate indirect interactions. Checkerboard analyses can determine if observational patterns in species co-occurrence differ from random assembly (Stone and Roberts 1990; Gotelli 2000). Such analyses, however, do not explicitly test for differences in interaction strengths, nor account for indirect interactions between species.

A new method using Markov networks can be a solution to these difficulties in estimating interaction strengths (D. J. Harris 2016). Markov networks are a promising method to get information about species interaction strengths while controlling for indirect interactions between species. Using this approach, pairwise interaction strengths are estimated from community observational data (D. J. Harris 2016). Here we define interaction strength as a measure of the degree of co-occurrence between pairs of species, akin to measuring the correlation between the occurrence of two species (Berlow et al. 2004). This method generates conditional species interaction strengths using maximum likelihood estimation from vectors of presences and absences. If species are less likely to occur together, their interaction strength will be negative. And conversely if species are more likely to occur together, their interaction strength will be positive. While this method was developed for competitive networks that show a “checkerboard” pattern, we argue that it is also useful to infer interaction strengths from certain types of simple food webs (see also Harris 2016). Even though it would seem that a predator cannot persist in the absence of its prey in a closed system, open systems with a high colonization rate of the prey and a high predation rate can also display a “checkerboard” pattern between the predator and the prey. When the predator consumes its prey to extinction, we may find the predator on its own. If the prey has a high colonization rate, it can colonize patches where the predator is absent. These colonization - extinction dynamics lead to a system with patch dynamics (e.g. Englund et al. 2009). In other words, the spatial scale can affect the degree of co-occurrence observed between predators and prey; at small scales, effective predators should reduce or eliminate their prey (negative co-occurrence) while at larger scales predators and prey should co-occur (Freilich et al. 2018).

Despite their potential, Markov methods have thus far not been used to reconstruct interactions in real food webs along environmental and spatial gradients. Good candidate ecosystems for such analyses are insular systems with simple food webs that occur over wide geographic areas. A classical food web model system that fits these criteria are the aquatic communities that live inside bromeliad plants in the neotropics; these communities often occur as clusters that exchange species via dispersal (i.e. metacommunities). In these communities, a suite of voracious predators can limit the abundance of prey species (Hammill et al. 2014), and prey colonizationis are rapid (Hammill et al. 2015). Thus bromeliad communities are suitable model systems to test how environment and space can affect species interactions and community composition (Petermann et al. 2015; Farjalla et al. 2012).

Making use of this model system, we explored three main questions. First, we tested whether environmental conditions varied between our sampling sites, located along a geographic gradient. Second, we describe change in community composition along this geographic gradient, and then partitioned this between-site variance in species composition into the effects of environment and distance. Third, we used Markov networks to quantify species interactions at each site. We explored whether difference between sites in the strength of species interactions could be explained by geographic variation in environmental conditions.

## Methods

### Model system

Tank bromeliads accumulate water inside their leaf axils, providing habitat for communities of aquatic macroinvertebrates (Kitching 2000). Inside each bromeliad, these aquatic macroinvertebrates interact to form a food web comprised of detritivores, filter feeders, intermediate predators and top predators. Bromeliad macroinvertebrate communities are known to be particularly sensitive to changes in precipitation, since this can change the amount of habitat available for the invertebrates (Pires et al. 2016). For example, drought in bromeliads is known to reduce growth rates of some invertebrate species (Amundrud and Srivastava 2015). Therefore, we expect that changes in precipitation have the potential to substantially affect species interactions and community composition.

### Study Area

The study area was located in the sand dunes of coastal Brazil (Figure 1), in the states of Rio de Janeiro and São Paulo. We sampled ten sites, seven of which were within the Jurubatiba National Park in Rio de Janeiro state, Brazil. The other three sites were located in the sand dunes of Arraial do Cabo (Rio de Janeiro), Marica (Rio de Janeiro), and Ilha Bela (Sao Paulo). This sampling design resulted in the sites closest to Jurutabita National Park receiving low precipitation, the sites close to Marica receiving intermediate precipitation, and the sites closest to Ilha Bela and Arraial do Cabo receiving high precipitation in the month immediately before sampling (February and March 2015, Appendix S1: Figure S1, S2).

**Figure 1.**
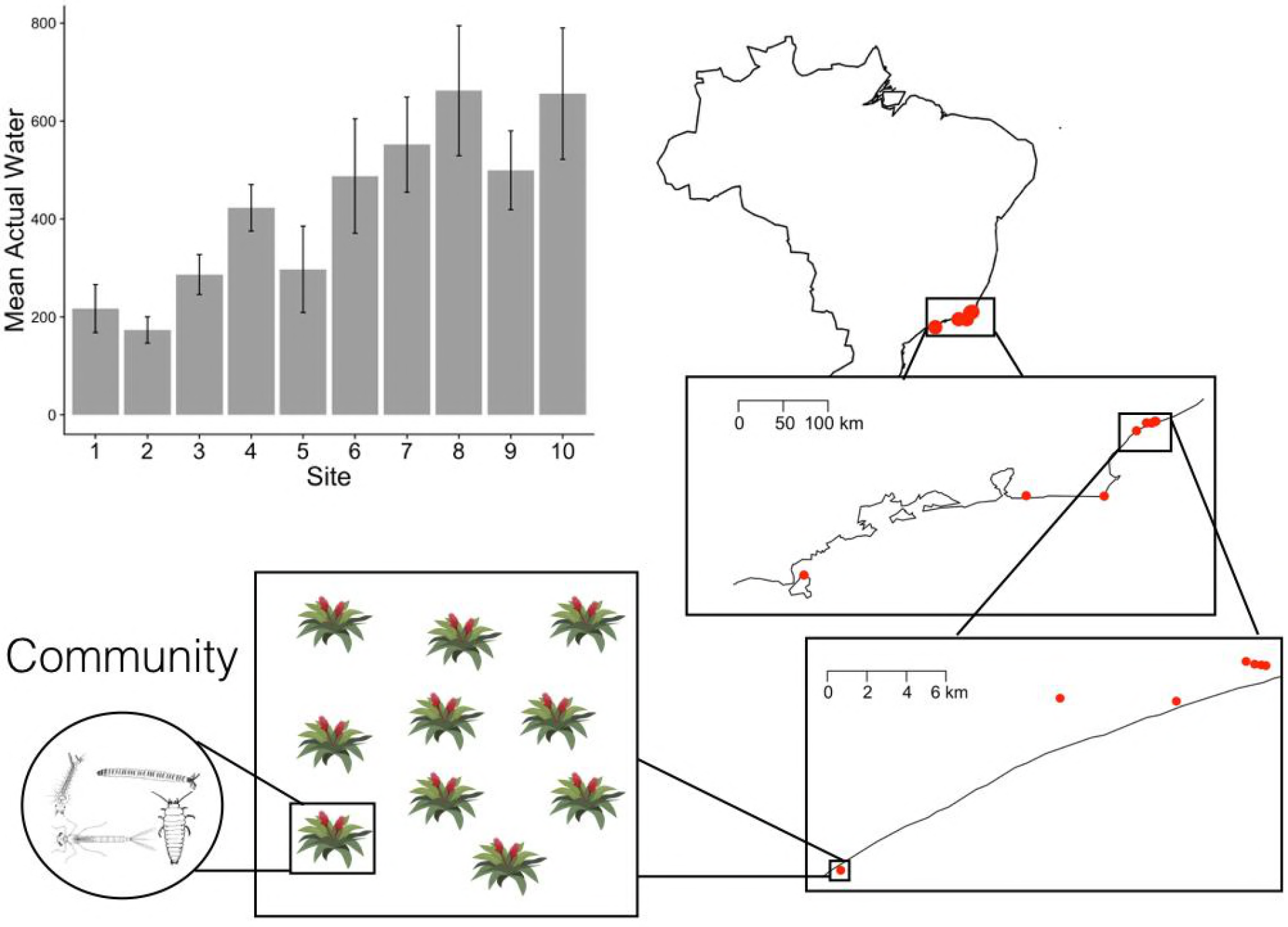
Locations of sites across the eastern coast of Brazil. A community is the set of species found in one bromeliad. A site is comprised of ten bromeliads found within 100 meters of each other. Ten sites were sampled, with a hierarchy of distances between bromeliads (nested boxes, right side of diagram). The mean actual water found in the bromeliads from each site is shown. Bars represent mean and standard error of the mean. Sites 1 to 10 are ordered north to south.

### Sampling

We sampled all macroinvertebrate communities between March and May 2015. In each metacommunity, we dissected ten bromeliads to collect all the invertebrates in each plant. The invertebrate samples were preserved in 99% ethanol. Macroinvertebrates were counted and identified to genus level whenever possible. For every bromeliad, we measured a suite of environmental variables to assess the amount and quality of habitat available to the invertebrates: the height (cm) and diameter (cm, measured as the maximum distance between leaf tips) of the plant, maximum water volume (mL, calculated by emptying the plant and calculating how much water the plant could hold before it overflowed), actual water volume (mL), longest leaf length (cm), longest leaf width (cm), number of leaves, canopy cover (% of shaded pixels in photos taken looking directly up from the bromeliad), total detritus (g dry mass), pH, oxygen concentration (% saturation), salinity (ppt), temperature (°C), and turbidity (NTU). Water chemistry and temperature variables were measured using a portable multiparameter waterproof meter in the field as soon as the water was collected from the plant.

## Data analysis

### Environmental variation between sites

We established if sites differed in their environmental conditions using simple ANOVA for most environmental variables. For oxygen saturation and canopy cover we used an analogous generalized linear model specifying a binomial family error distribution as is suitable for percentage data restricted between 0 and 100 (Appendix S1).

### Compositional variation between sites

We tested for differences in community composition between sites using permutational multivariate analysis of variance using distance matrices (function *adonis* in R package *vegan*, hereafter referred to as adonis). Multivariate tests of dispersion (function *betadisper* in R package *vegan*) were used to test for differences in community variation (beta diversity) between geographic sites. We summarized abundances according to genus so that these results would be comparable with species interactions analyses. For the *adonis* analysis, we tested if bromeliads from different sites and containing different water volumes had different average differences in community composition. We used water volume in this analysis since, of all the environmental variables, it differed the most between sites. For the multivariate test of dispersion, we tested if bromeliads from different sites differed in their beta diversity, where within-site beta diversity was measured as the average dissimilarity of bromeliad invertebrate communities from the centroid in multivariate space (Anderson 2001, 2006). To visualize these results, we represented the differences between bromeliads in their community composition with non-metric multidimensional scaling (NMDS) plots (Anderson 2001). This type of plot shows both the differences between sites in their average community composition (position of centroids) as well as differences between sites in beta diversity (the standardised residuals around the centroids).

### Species Interactions

To obtain species interactions strengths, we used Markov network analysis (D. J. Harris 2016). This method does not make any assumptions about the topology of the food web, nor do we have to define which species might interact with each other. The analysis calculates conditional species interaction strength using maximum likelihood estimation from presence/absence data. We summarized abundances according to genus, to reduce computational complexity. The trophic role of bromeliad aquatic invertebrates is highly conserved at the genus level (Poff et al. 2006), so we likely have not averaged over different trophic interactions with this approximation. The abundance data of each genus were transformed into presence/absence data. We performed Markov Network analysis separately for each site (D. J. Harris 2016). The output of this analysis is the relative interaction strength for every pair of species in the site. We define these interaction strengths as relative because all interactions occurring within a site are drawn from a logistic distribution density function. Note that the prior distribution determines the final distribution of the interaction strengths.

### Confirmation of suitability of Markov network method

We first wanted to confirm if the Markov network method was able to recover known interaction strengths and could predict trophic interaction strengths in simple bromeliad food webs. We took two different approaches to this confirmations. First, we ran the Markov network analysis on a three species module from Costa Rica where all interaction strengths had been established based on experiments (Hammill, Atwood, and Srivastava 2015). We were able to get similar interaction strengths as those expected based on direct experimentation (Appendix S2, Table S1). A more detailed discussion of the suitability and limitations of the methods is provided in Appendix S3.

Second, because we have prior knowledge on the trophic ranks of every genera in the Brazilian dataset, we could test whether the Markov network method could correctly assign the trophic positions of genera. For this analysis, we started by assessing the types and strengths of interactions for every combination of genera. Positive interactions mean that species are more likely to co-occur than expected by chance. Negative interactions mean that species are less likely to co-occur than expected by chance. We calculated the number of positive and negative interactions for every species, regardless of the strength across sites; all values above zero are positive and all values below zero are negative. We classified every species as a predator or a prey, to test if known differences in trophic positions have a different preponderance of positive (expected for prey) or negative (expected for predators) interactions. The species or genera known to be predatory are: the damselfly *Leptagrion andromache* nymphs, elephant mosquito larvae *Toxorynchites*, *Corethrella* midge larvae, horsefly larvae *Tabanidae* and cranefly larvae *Tipulidae*. For this analysis, we explained the total number of interactions for each genera as a function of the sign of the interaction (either positive or negative) and the trophic position. Since individual genera are present in different interactions, we included genera identity as a random effect in a generalized mixed effect model. The independent units of replication are the sites. We used a Poisson error distribution, appropriate for left-skewed count data. The interaction term of this model, between trophic position and the sign of the between-genera interaction, tests if predators and prey differ in the sign of their biological interactions.

### Effect of environment on species interactions

Once we were able to confirm that Markov Network analysis correctly distinguished between predators and prey in terms of the predominant sign of interactions, we could then examine if the environment explained differences between metacommunities in the relative strength of either positive or negative interactions. For this analysis, we separated negative from positive interactions to assess how interaction strength (within a particular sign) changes with the environmental variables, based on linear and quantile regressions. Quantile regressions are useful when there is unequal variation in the data and therefore there might be more than one slope describing the relationship between response variable and predictor. Quantile regression is also more robust to outliers than mean regressions (Cade and Noon 2003). The linear regression and quantile regression p values were adjusted using the Holm correction for multiple comparisons. To confirm the robustness of our results, we performed permutation analysis shuffling community composition. To further understand our results, we partitioned beta diversity between nestedness and turnover (See Appendix S5 for further details).

For the species interaction analyses we used the *rosalia* package (D. Harris 2015), all multivariate analyses were performed using the *vegan* package (Oksanen et al. 2017), mixed effect models were performed using *lme4* (Bates et al. 2015) and *car* (Fox and Weisberg 2011), and all analyses were done using the R programming language (R Core Team 2016).

## Results

### Environmental variation between sites

The only two environmental variables that significantly differed between metacommunities were maximum and actual water volume in bromeliads (Appendix S1: Table S1), and of these two, the most pronounced gradient was observed in the actual water volume in the bromeliads (F_10, 90_ = −3.854, P = 0.0003, Figure 1). We therefore focus on actual water volume as the major environmental variable for analyses with univariate explanatory variables.

### Community variation along an environmental gradient

Community composition differed between sites depending on the actual water volume in the bromeliads (F_9, 90_ = 4.649, P = 0.001, Figure 2a, b), however, beta diversity, measured as multivariate dispersion around site centroids, differed only marginally among sites (F_9, 90_ = 1.966, P = 0.052). These site differences in beta diversity were mainly driven by the sites that were the furthest apart geographically and differed the most in the actual water contained in bromeliads (post-hoc pairwise adonis and tukey’s tests: Appendix S5: Table S1). Using variation partitioning, we confirmed that the majority of the variation in community composition between bromeliads was explained by the partial (i.e. pure) effects of environmental variables (20%: F_11, 76_ = 3.515, P = 0.001) and less so by spatial variables (14%: F_5, 76_ = 4.641, P = 0.001), with only a small amount (5%) due to the covariance between environment and the space (Appendix S2, Figure S1). The difference in community composition between sites was mostly due to species turnover (70%) and not due to nestedness (30%, Figure 2c). Therefore species were not being progressively lost along the gradient of actual water in the bromeliads and specie richness per bromeliad was relatively constant across the gradient (Figure 2a).

**Figure 2.**
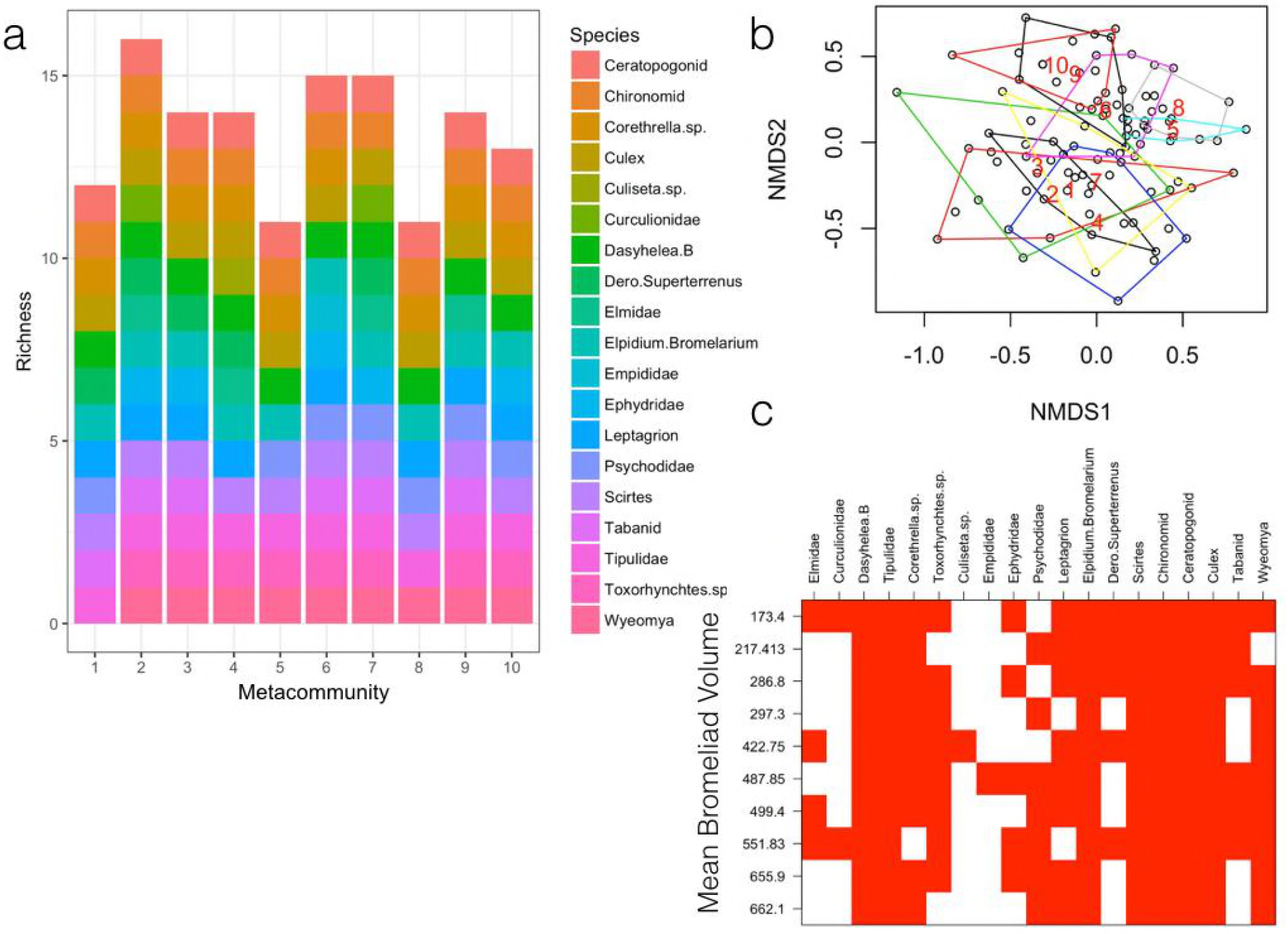
Comparison of the community composition between sites. A. Shows the different species present in each site. B. The community composition of every bromeliad is compressed into two axes. Each site is represented by a polygon containing all bromeliads within that site. The stress of the NMDS plot was 0.26. C. A species (columns) by sites (rows) matrix with sites ordered based on mean water volume, showing that the differences in community composition across water volume are due to species turnover and not nestedness of losing species along that gradient.

### Confirmation of Markov network applicability to trophic interactions

We first confirmed that the Markov network method could correctly identify trophic interactions in our dataset. We calculated the strength of species interactions for every site (Figure 3). The top predator *Leptagrion andromache* dominated negative interactions (Figure 4) which is expected since it preys on most species in the community. In general, prey species had more positive interactions and predator species had more negative interactions compared to random expectations (β = 0.564, z value = 4.456, P value = 8.3 × 10 ^− 6^, Figure 4). This result was robust to the matrix permutations of presences (P value = 0, Figure A3).

**Figure 3:**
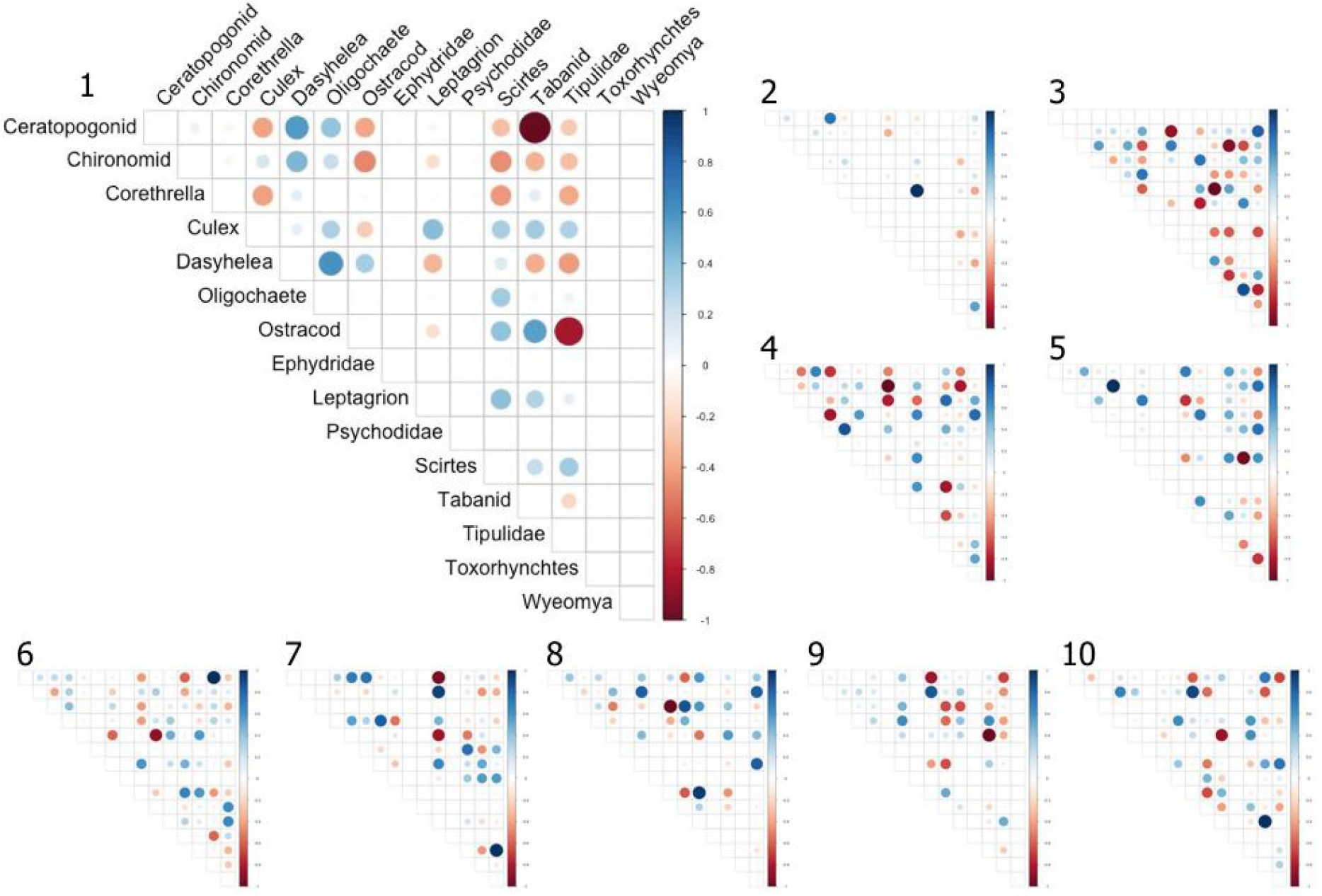
Relative strength of species interactions in every site. Species interactions are scaled to 1. Where rows or columns are empty, that particular species is not in that site.

**Figure 4:**
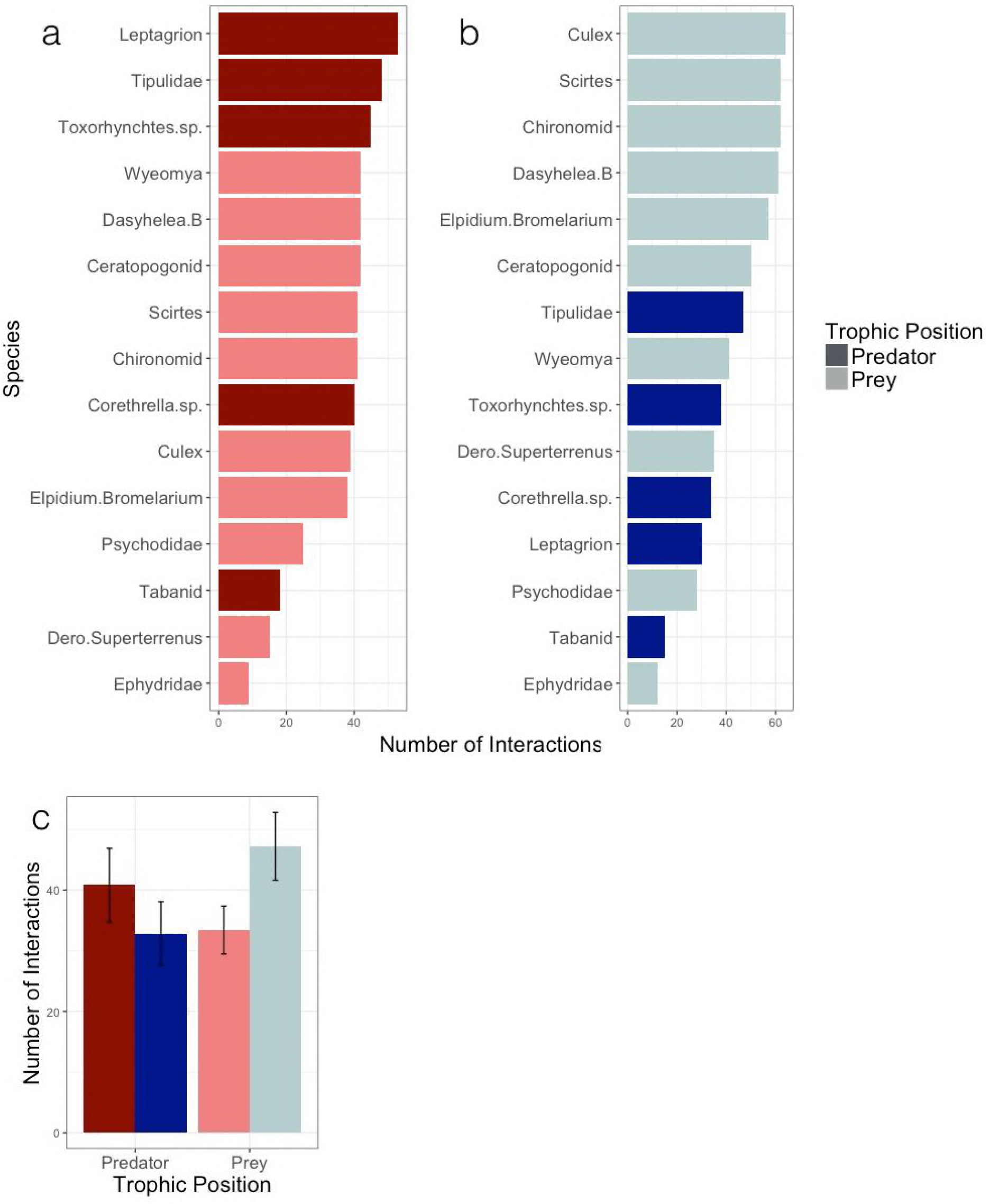
Compiled number of a) negative and b) positive interactions for each species across all metacommunities. Darker colours indicate known predatory species prior to the analysis and lighter colours indicate prey species. c) The total number of interactions for different trophic levels and the type of interaction. Blue indicates positive interactions and red indicate negative interactions. Predators have a higher number of negative interactions while prey have a higher number of positive interactions. Bars represent mean and standard error of the mean.

### Effect of environment on species interactions

As the majority of genera were found in most sites, we could ask how each genus differed across large scale environmental gradients in the type (interaction sign) and interaction strength with other community members. Using site means of actual water as the environmental gradient, we found that the relative strength of positive and negative interactions remain constant between sites for most genera, but for *Tipulidae*, *Wyeomyia* and *Elpidium bromeliarium* the relative strength of negative interactions diminished with site water volume. That is, sites whose bromeliads contained less water on average tended to have stronger negative interactions between *Tipulidae* (linear regression: β = 1.179 × 10^−3^, P value = 0.017), *Wyeomyia* (β = 8.254 × 10^−4^, P value = 0.062) and *Elpidium bromeliarum* (β = 1.257 × 10^−3^, P value = 0.016) and individuals and other community members. We also explored Box-Cox transformations but the results did not vary qualitatively. Arguably, quantile regression might be better suited to detecting changes in the distribution of interactions, in which case only the *Tipulidae* negative interactions are still related to the mean water volume, even after the results were adjusted for multiple comparisons (first quantile regression: β = 1.151 × 10 ^−3^, P value = 0.05, Figure 5a).

**Figure 5:**
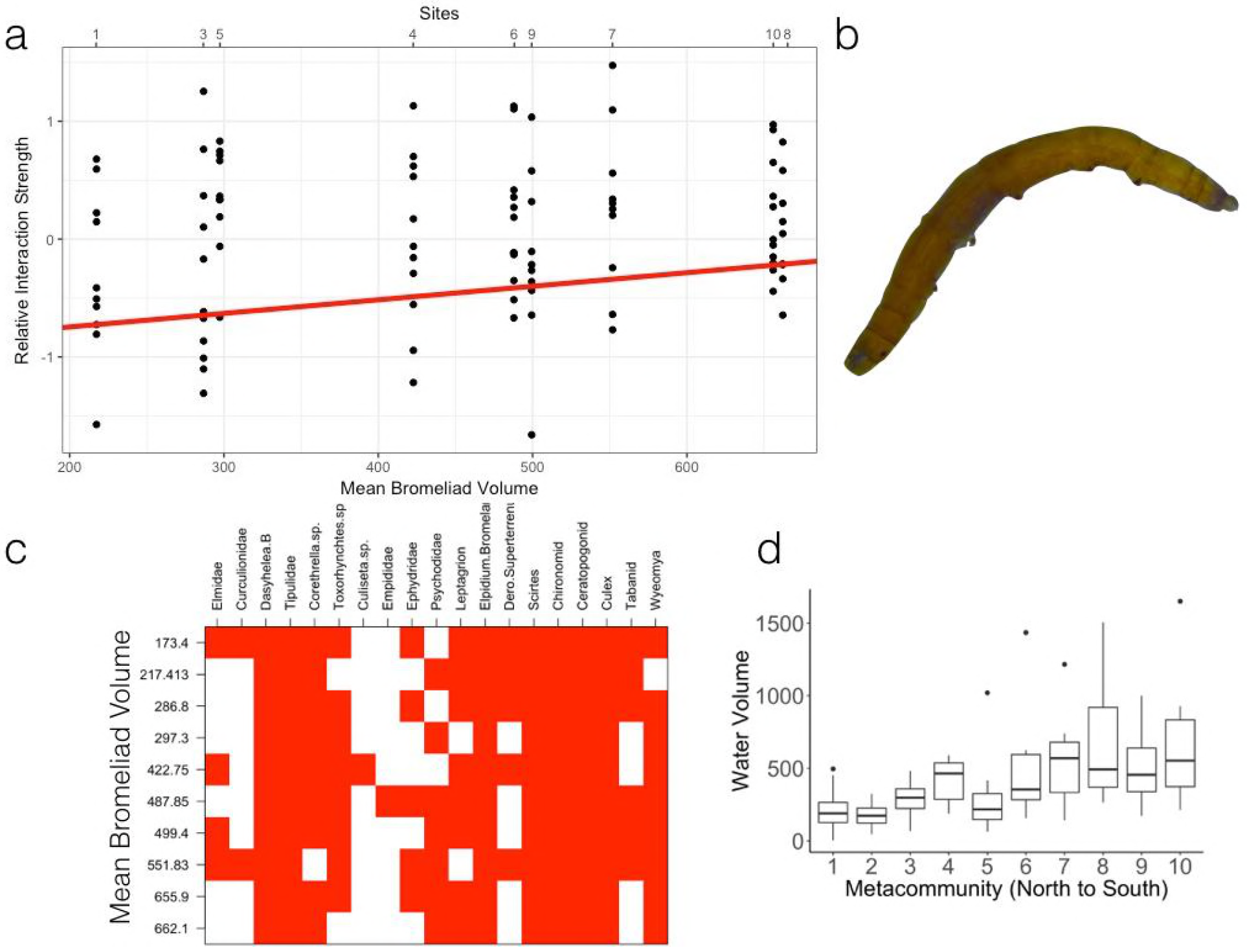
A. The relative interaction strength of tipulids as a function of the mean water volume of the metacommunity. The red line represents the first quantile regression. B. Image of a Tipulid larvae.

Even though difference between sites in interactions strengths could be due either to changes in the per capita interaction strengths between specific taxa, or changes in the pool of species available for interactions, we find that low volume sites do not progressively lose species (i.e. there is no nested loss of species, Figure 2a). We also found that the absence of a species due to low water volume was not related to the interaction strength between those species and the tipulid (Appendix S2, Figure S6)

## Discussion

The main conclusions of this study were threefold. First, we explored the main axes of environmental variation that could separate our sampled sites. We found that, due to variation in precipitation, the actual water contained in the bromeliads was the main variable that consistently varied between sites. Second, we found that gamma diversity varied between sites. However, most of that variation was due to the turnover of some species and not due to sequential loss of species being filtered by the environmental gradient (i.e. higher turnover vs. nestedness). In addition, we found that more variation in community composition could be attributed to the environment than to space. Third we used a Markov network approach to quantify trophic interactions at the site level and used this tool to explore how environmental conditions at the site level are linked to geographic variation in the strength of trophic interactions. We first confirmed that the results of the Markov Network analysis matches our knowledge of natural history of the system, finding that known predators are more often engaged in negative interactions. We then showed that the interactions between tipulids and other species changed along a site gradient in bromeliad water volume; sites with lower water volume had more intense negative interactions.

Our sites were located along a precipitation gradient with those in the south-west of our gradient receiving less rainfall than those towards the north-east (Appendix S1: Figure S1, S2). This gradient was reflected in the amount of actual water found in the bromeliads. Even though it seems that these sites were otherwise similar, low water volumes can affect bromeliad networks through a multitude of mechanisms: (i) low water volumes can select species whose traits allow them to be tolerant to drought (Dézerald et al. 2015); (ii) low water volumes decrease habitat size thereby decreasing the size of the ecological network (Dézerald et al. 2013) (iii) high water volumes allow higher trophic levels to persist in the local network (Amundrud and Srivastava 2015).

Overall, community composition differed along the geographic gradient in bromeliad water volume. However, this difference is not driven by the sequential loss of species along this gradient, but instead turnover in species identity. For example, the oligochaete *Dero superterrenus* was more common in the drier sites, and *Polypedilum* chironomids in the wetter sites. Such turnover may be related to the life history of organisms: oligochaetes reproduce within bromeliads, and so are resident year round, whereas larval chironomids require terrestrial adults to oviposit eggs and adults may delay oviposition until most bromeliads in the site are water-filled. Previous studies have shown that the functional traits of bromeliad invertebrates determine their response to water levels within bromeliads: taxa able to survive low water conditions are characterized by small size and deposit or filter feeding whereas taxa able to rapidly colonize full bromeliads are characterized by drought-tolerant eggs and short generation times (Dézerald et al. 2015). Since species traits determine their response to altered environmental conditions, selection of species through their traits can alter not only the size of a network but also its architecture (Tylianakis and Morris 2017a). More generally, if species differ in their optimal environment due to their life history and tolerance traits, we would expect that the arrangement of sites along an environmental gradient would cause a turnover in species composition due to species sorting mechanisms or when early successional species are gradually lost (Mathew A. Leibold and Chase 2017; Brendonck et al. 2014). Note that this species turnover occurs despite constant species richness between sites, regardless of water volume. However, within site, species richness increases with bromeliad water volume, as shown in previous studies of bromeliad invertebrates (Jabiol et al. 2009).

Even though Markov network analysis can be used in trophic networks, the low degrees of freedom in compositional data only allow us to estimate one interaction value per species pair (D. J. Harris 2016). However, we can use information from the natural history of the system to allow us to interpret these interactions (D. J. Harris 2016; Freilich et al. 2018). We looked at the type of interactions that prey and predators participated in, knowing prior to the analysis which species were predators and which species were prey. We found that the top predator *Leptagrion andromache* dominates negative interactions, as expected from a generalist predator known to have high per capita impact on its prey (Hammill, Atwood, and Srivastava 2015). Furthermore, we found that predatory species were more likely to participate in net negative interactions and prey species were more likely to participate in net positive interactions, confirming that the Markov network approach could detect trophic interactions. This however, does not mean that all predator-prey relationships are necessarily detected via negative interaction strengths.

Our Markov analyses indicated that, while overall species interactions are similar in sign and strength along the water volume gradient, for three genera there is a consistent pattern of strengthening negative interactions in sites with lower water level. This pattern was particularly robust for tipulids. There are two possible mechanisms for this result. First, species that have only weak interactions with the three genera may become absent at sites with low water volumes, allowing stronger negative interactions to influence the mean. If this mechanism was operating, then the pattern should disappear with quantile regression. Indeed for two genera it does, but not for tipulids. The second mechanism is that many of the negative interactions intensify in strength as site water volumes diminish. This mechanism is consistent with the patterns seen in the tipulids.

Tipulids may show stronger negative interactions at low water volumes because they become generalist predators. This mechanism is supported by previous research, which found that tipulids in Costa Rican bromeliads supplement detritivory with opportunistic predation under drought (Amundrud and Srivastava 2016). These authors hypothesized that decreasing water volume in a bromeliad restricts the space for prey movement, and therefore the tipulids can become more effective predators. Other manipulative experiments confirm that bromeliad predators are more effective in smaller water volumes (Diane S. Srivastava 2006). Generally, from a biomechanical perspective, consumption rates of predators should depend on habitat dimensionality because it influences the cost of locomotion and the probability of prey escape (Pawar, Dell, and Savage 2012).

Tipulids thus appear to be facultative predators, opportunistically switching from detritivory to predation. Facultative predators feed both on plant matter and animals at the same developmental stage; they represent a case of non-obligate omnivory (Albajes and Alomar 2004). Facultative predation may constitute an adaptive strategy in habitats with high variability of food sources, and allow species to withstand changing environments (D. S. Srivastava et al. 2008; Albajes and Alomar 2004). Bromeliad habitats are known to be very variable, with water levels that fluctuate year around (Scarano 2002). Therefore facultative predation may be a favourable strategy in these systems.

Previous studies of bromeliads infauna corroborate our finding that trophic interactions may change with climate. For example, over a much larger geographical gradient, Romero et al. (2016) found that cooler, less seasonal climatic conditions resulted in stronger top-down control from predators, based on biomass ratios of top predators to detritivores as a proxy for interaction strength. Although this result may appear to be the opposite to our finding that tipulid predation intensified in warmer, more seasonal sites, we note that the top predators in the Romero et al. study are dominated by odonate larvae. Tipulids have until recently not been considered predators in bromeliad food webs, and so were considered detritivores in Romero et al. (2016), but they could be classified as mesopredators. There was no effect of climatic stability on the other meso-predators in (Romero et al. 2016). An intriguing topic for future study is whether seasonal droughts in Rio de Janeiro state, Brazil, shift predation from odonates to tipulids.

Our study reinforces the general point that networks can change along an environmental gradient through two main ways (i) through the turnover of species or (ii) through the change in species interactions (Tylianakis and Morris 2017b). Here we found both mechanisms are contributing to the changes in community composition along a gradient of actual water in bromeliads.

## Conclusion

In this study we provided evidence for changes in network structure along an environmental gradient through two mechanisms. First we showed that community composition differed along a gradient of actual water in macroinvertebrate communities due to the turnover of some species. Second we showed that species interactions also differed along this gradient. In our system, lower water levels likely changed the effectiveness of different predation strategies reflected in different more negative species interactions. The notion that the same actors might be active in a totally different play, implies that it may be not be recommendable to directly link species composition to ecosystem functioning as attempted in many recent studies. Broader applications of the Markov approach to assess interaction strengths could assist studies that aim to explain differences in functional aspects of ecosystems that cannot be attributed to differences in species composition.

## Acknowledgements

This work was partially funded by CNPq through PVE grants (Process 400454/2014-9) and the Mitacs Globalink scholarships. L.M.G. is supported by NSERC CGS-D and UBC Four Year Fellowships. D.S.S is supported by NSERC Discovery Grants. V.F.Farjalla is partially supported by a CNPq productivity grant.

## References

Albajes, R., and O. Alomar. 2004. “Facultative Predators.” In Encyclopedia of Entomology, 818–23.

Amundrud, Sarah L., and Diane S. Srivastava. 2015. “Drought Sensitivity Predicts Habitat Size Sensitivity in an Aquatic Ecosystem.” Ecology 96 (7): 1957–65.

Amundrud, Sarah L., and Diane S. Srivastava. 2016. “Trophic Interactions Determine the Effects of Drought on an Aquatic Ecosystem.” Ecology 97 (6): 1475–83.

Anderson, Marti J. 2001. “A New Method for Non-Parametric Multivariate Analysis of Variance.” Austral Ecology 26 (1): 32–46.

Anderson, Marti J.. 2006. “Distance-Based Tests for Homogeneity of Multivariate Dispersions.” Biometrics 62 (1): 245–53.

Bates, Douglas., Martin Maechler, Ben Bolker, Steve Walker (2015). Fitting Linear Mixed-Effects Models Using lme4. Journal of Statistical Software, 67(1), 1–48.doi:10.18637/jss.v067.i01

Berlow, Eric L., Anje-Margiet Neutel, Joel E. Cohen, Peter C. de Ruiter, Bo Ebenman, Mark Emmerson, Jeremy W. Fox, et al. 2004. “Interaction Strengths in Food Webs: Issues and Opportunities.” The Journal of Animal Ecology 73 (3): 585–98.

Blanchet, F. Guillaume, Pierre Legendre, and Daniel Borcard. 2008. “Forward Selection of Explanatory Variables.” Ecology 89 (9): 2623–32.

Borcard, Daniel, Francois Gillet, and Pierre Legendre. 2011. Numerical Ecology with R. Springer.

Brendonck, Luc, Merlijn Jocqué, Karen Tuytens, Brian V. Timms, and Bram Vanschoenwinkel. 2014. “Hydrological Stability Drives Both Local and Regional Diversity Patterns in Rock Pool Metacommunities.” Oikos 124 (6): 741–49.

Cade, Brian S., and Barry R. Noon. 2003. “A Gentle Introduction to Quantile Regression for Ecologists.” Frontiers in Ecology and the Environment 1 (8): 412.

Connolly, J., and P. Wayne. 1996. “Asymmetric Competition between Plant Species.” Oecologia 108 (2): 311–20.

Dézerald, Olivier, Régis Céréghino, Bruno Corbara, Alain Dejean, and Céline Leroy. 2015. “Functional Trait Responses of Aquatic Macroinvertebrates to Simulated Drought in a Neotropical Bromeliad Ecosystem.” Freshwater Biology 60 (9): 1917–29.

Dézerald, Olivier, Stanislas Talaga, Céline Leroy, Jean-François Carrias, Bruno Corbara, Alain Dejean, and Régis Céréghino. 2013. “Environmental Determinants of Macroinvertebrate Diversity in Small Water Bodies: Insights from Tank-Bromeliads.” Hydrobiologia 723 (1): 77–86.

Edgerly, J. S., M. S. Willey, and T. Livdahl. 1999. “Intraguild Predation among Larval Treehole Mosquitoes, Aedes Albopictus, Ae. Aegypti, and Ae. Triseriatus (Diptera: Culicidae), in Laboratory Microcosms.” Journal of Medical Entomology 36 (3): 394–99.

Englund, Göran, Frank Johansson, Patrik Olofsson, Juha Salonsaari, and Johanna Ohman. 2009. “Predation Leads to Assembly Rules in Fragmented Fish Communities.” Ecology Letters 12 (7): 663–71.

Farjalla, Vinicius F., Diane S. Srivastava, Nicholas A. C. Marino, Fernanda D. Azevedo, Viviane Dib, Paloma M. Lopes, Alexandre S. Rosado, Reinaldo L. Bozelli, and Francisco A. Esteves. 2012. “Ecological Determinism Increases with Organism Size.” Ecology 93 (7): 1752–59.

Fincke, Ola M. 1994. “Population Regulation of a Tropical Damselfly in the Larval Stage by Food Limitation, Cannibalism, Intraguild Predation and Habitat Drying.” Oecologia 100 (1-2): 118–27.

Fox, John. and Sanford Weisberg (2011). An {R} Companion to Applied Regression, Second Edition. Thousand Oaks CA: Sage. URL: http://socserv.socsci.mcmaster.ca/jfox/Books/Companion

Freilich, Mara A., Evie Wieters, Bernardo R. Broitman, Pablo A. Marquet, and Sergio A. Navarrete. 2018. “Species Co-Occurrence Networks: Can They Reveal Trophic and Non-Trophic Interactions in Ecological Communities?” Ecology 99 (3): 690–99.

Gotelli, Nicholas J. 2000. “Null Model Analysis of Species Co-Occurrence Patterns.” Ecology 81 (9): 2606.

Hammill, Edd, Trisha B. Atwood, Paloma Corvalan, and Diane S. Srivastava. 2014. “Behavioural Responses to Predation May Explain Shifts in Community Structure.” Freshwater Biology 60 (1): 125–35.

Hammill, Edd, Trisha B. Atwood, and Diane S. Srivastava. 2015. “Predation Threat Alters Composition and Functioning of Bromeliad Ecosystems.” Ecosystems 18 (5):857–66.

Harris, David. 2015. Rosalia: Exact Inference for Small Binary Markov Networks. R Package (version 0.1.0). Http://dx.doi.org/10.5281/zenodo.17808.

Harris, David J. 2016. “Inferring Species Interactions from Co-Occurrence Data with Markov Networks.” Ecology 97 (12): 3308–14.

Jabiol, J., B. Corbara, A. Dejean, and R. Céréghino. 2009. “Structure of Aquatic Insect Communities in Tank-Bromeliads in a East-Amazonian Rainforest in French Guiana.” Forest Ecology and Management 257 (1): 351–60.

Kitching, R. L. 2000. Food Webs and Container Habitats: The Natural History and Ecology of Phytotelmata. Cambridge University Press.

Kraft, Nathan J. B., Peter B. Adler, Oscar Godoy, Emily C. James, Steve Fuller, and Jonathan M. Levine. 2014. “Community Assembly, Coexistence and the Environmental Filtering Metaphor.” Functional Ecology 29 (5): 592–99.

Laliberté, Etienne, and Jason M. Tylianakis. 2010. “Deforestation Homogenizes Tropical Parasitoid-host Networks.” Ecology 91 (6): 1740–47.

Leibold, M. A., M. Holyoak, N. Mouquet, P. Amarasekare, J. M. Chase, M. F. Hoopes, R. D. Holt, et al. 2004. “The Metacommunity Concept: A Framework for Multi-Scale Community Ecology.” Ecology Letters 7 (7): 601–13.

Leibold, Mathew A., and Jonathan M. Chase. 2017. Metacommunity Ecology. Princeton University Press.

Maser, Gabriel L., Frédéric Guichard, and Kevin S. McCann. 2007. “Weak Trophic Interactions and the Balance of Enriched Metacommunities.” Journal of Theoretical Biology 247 (2): 337–45.

Oksanen, Jari, Guillaume Blanchet, Michael Friendly, Roeland Kindt, Pierre Legendre, Dan McGlinn, Peter Minchin, et al. 2017. Vegan: Community Ecology Package (version 2.4-2). https://cran.r-project.org/web/packages/vegan/.

Paine, Robert T. 1966. “Food Web Complexity and Species Diversity.” The American Naturalist 100 (910): 65–75.

Pawar, Samraat, Anthony I. Dell, and Van M. Savage. 2012. “Dimensionality of Consumer Search Space Drives Trophic Interaction Strengths.” Nature 486 (7404): 485–89.

Petermann, Jana S., Vinicius F. Farjalla, Merlijn Jocque, Pavel Kratina, A. Andrew M. MacDonald, Nicholas A. C. Marino, Paula M. De Omena, et al. 2015. “Dominant Predators Mediate the Impact of Habitat Size on Trophic Structure in Bromeliad Invertebrate Communities.” Ecology 96 (2): 428–39.

Pires, Aliny P. F., Nicholas A. C. Marino, Diane S. Srivastava, and Vinicius F. Farjalla. 2016. “Predicted Rainfall Changes Disrupt Trophic Interactions in a Tropical Aquatic Ecosystem.” Ecology 97 (10): 2750–59.

Poff, N. Leroy, N. LeRoy Poff, Julian D. Olden, Nicole K. M. Vieira, Debra S. Finn, Mark P. Simmons, and Boris C. Kondratieff. 2006. “Functional Trait Niches of North American Lotic Insects: Traits-Based Ecological Applications in Light of Phylogenetic Relationships.” Journal of the North American Benthological Society 25 (4): 730–55.

Rall, Björn C., Ulrich Brose, Martin Hartvig, Gregor Kalinkat, Florian Schwarzmüller, Olivera Vucic-Pestic, and Owen L. Petchey. 2012. “Universal Temperature and Body-Mass Scaling of Feeding Rates.” Philosophical Transactions of the Royal Society of London. Series B, Biological Sciences 367 (1605): 2923–34.

R Core Team. 2016. R: A Language and Environment for Statistical Computing. R Foundation for Statistical Computing, Vienna, Austria (version 3.3.2). https://www.r-project.org/.

Romero, Gustavo Q., Gustavo C. O. Piccoli, Paula M. de Omena, and Thiago Gonçalves-Souza. 2016. “Food Web Structure Shaped by Habitat Size and Climate across a Latitudinal Gradient.” Ecology 97 (10): 2705–15.

Scarano, Fabio R. 2002. “Structure, Function and Floristic Relationships of Plant Communities in Stressful Habitats Marginal to the Brazilian Atlantic Rainforest.” Annals of Botany 90 (4): 517–24.

Srivastava, Diane S. 2006. “Habitat Structure, Trophic Structure and Ecosystem Function: Interactive Effects in a Bromeliad-insect Community.” Oecologia 149 (3): 493–504.

Srivastava, D. S., M. K. Trzcinski, B. A. Richardson, and B. Gilbert. 2008. “Why Are Predators More Sensitive to Habitat Size than Their Prey? Insights from Bromeliad Insect Food Webs.” The American Naturalist 172 (6): 761–71.

Stone, Lewi, and Alan Roberts. 1990. “The Checkerboard Score and Species Distributions.” Oecologia 85 (1): 74–79.

Tylianakis, Jason M., and Rebecca J. Morris. 2017a. “Ecological Networks Across Environmental Gradients.” Annual Review of Ecology, Evolution, and Systematics 48 (1): 25–48.

Tylianakis, Jason M., and Rebecca J. Morris. 2017b. “Ecological Networks Across Environmental Gradients.” Annual Review of Ecology, Evolution, and Systematics 48 (1): 25–48.

Wootton, J. Timothy. 1997. “Estimates and Tests of Per Capita Interaction Strength: Diet, Abundance, and Impact of Intertidally Foraging Birds.” Ecological Monographs 67 (1): 45.

